# Comparative Analysis of Association Networks Using Single-Cell RNA Sequencing Data Reveals Perturbation-Relevant Gene Signatures

**DOI:** 10.1101/2023.09.11.556872

**Authors:** Nima Nouri, Giorgio Gaglia, Hamid Mattoo, Emanuele de Rinaldis, Virginia Savova

## Abstract

Single-cell RNA sequencing (scRNA-seq) data has elevated our understanding of systemic perturbations to organismal physiology at the individual cell level. However, despite the rich information content of scRNA-seq data, the relevance of genes to a perturbation is still commonly assessed through differential expression analysis. This approach provides a one-dimensional perspective of the transcriptomic landscape, risking the oversight of tightly controlled genes characterized by modest changes in expression but with profound downstream effects. We present GENIX (Gene Expression Network Importance eXamination), a novel platform for constructing gene association networks, equipped with an innovative network-based comparative model to uncover condition-relevant genes. To demonstrate the effectiveness of GENIX, we analyze influenza vaccine-induced immune responses in peripheral blood mononuclear cells (PBMCs) collected from recovered COVID-19 patients, shedding light on the mechanistic underpinnings of gender differences. Our methodology offers a promising avenue to identify genes relevant to perturbation responses in biological systems, expanding the scope of response signature discovery beyond differential gene expression analysis.

**HIGHLIGHTS:**

- Conventional methods used to identify perturbation-relevant genes in scRNA-seq data rely on differential expression analysis, susceptible to overlooking essential genes.
- GENIX leverages cell-type-specific inferred gene association networks to identify condition-relevant genes and gene programs, irrespective of their specific expression alterations.
- GENIX provides insight into the gene-regulatory response to the influenza vaccine in naïve and recovered COVID-19 patients, expanding on previously observed gender-specific differences.

**GRAPHICAL ABSTRACT:** 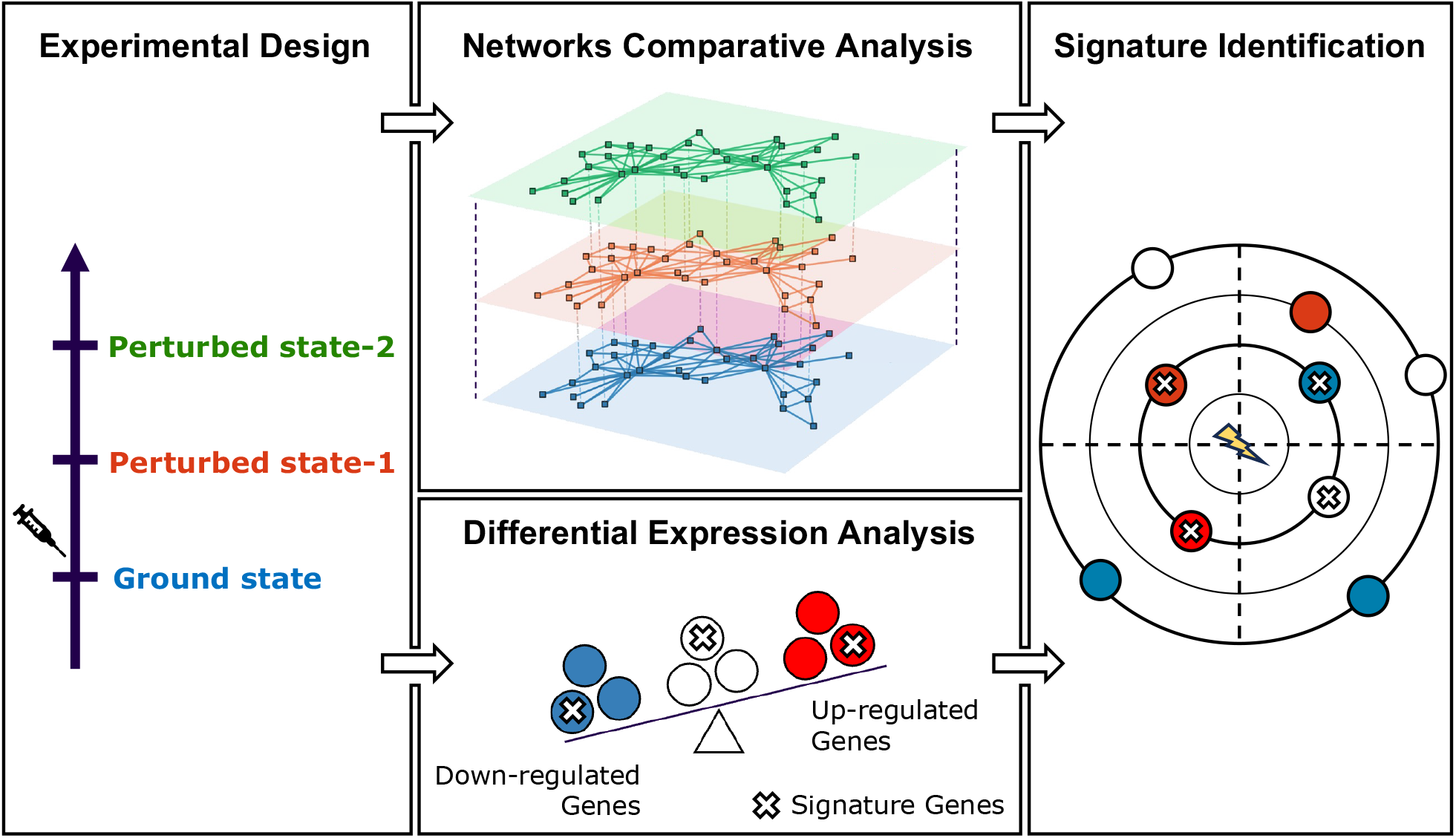

## INTRODUCTION

Identifying and prioritizing disease-associated or therapeutic response-related genes from a vast pool of candidates poses a significant challenge in modern drug discovery pipelines (Mohs, 2017; Jenwitheesuk, 2008). With 5-7 thousand genes, out of the approximately 20,000 protein-coding genes in the human genome, coexisting within cells at any given moment, data-driven strategies are required to discern the genes responsible for intra-and extra-cellular functions. Recent advancements in single-cell RNA sequencing (scRNA-seq) technology have revolutionized our ability to unravel the complex landscape of gene expression within tissues and organisms at the single-cell level (Yofe, 2020; Trombetta, 2014; Svensson, 2018; Stuart, 2019; Stubbington, 2017; Regev, 2017). This cutting-edge technology provides powerful means to comprehensively profile cell-level transcriptomic patterns, thereby facilitating the identification of crucial gene signatures associated with disease states and therapeutic responses (Van de Sande, 2023).

Differential expression analysis (DEA) is a widely employed method in transcriptomics research to identify gene expression alterations across different conditions or experimental groups (**Figure 1A**). This approach is invaluable, but has limitations. Notably, DEA is susceptible to overlooking genes that do not exhibit pronounced expression changes, which can be magnified downstream (e.g., transcription factors), or genes that undergo contextual changes in their interactions rather than changes in expression (e.g., signaling molecules) (**Figure 1B**). Therefore, relying exclusively on DEA may inadvertently exclude pivotal genes that play substantial roles in the experimental or biological condition despite subtle transcription level changes.

**Figure 1:**
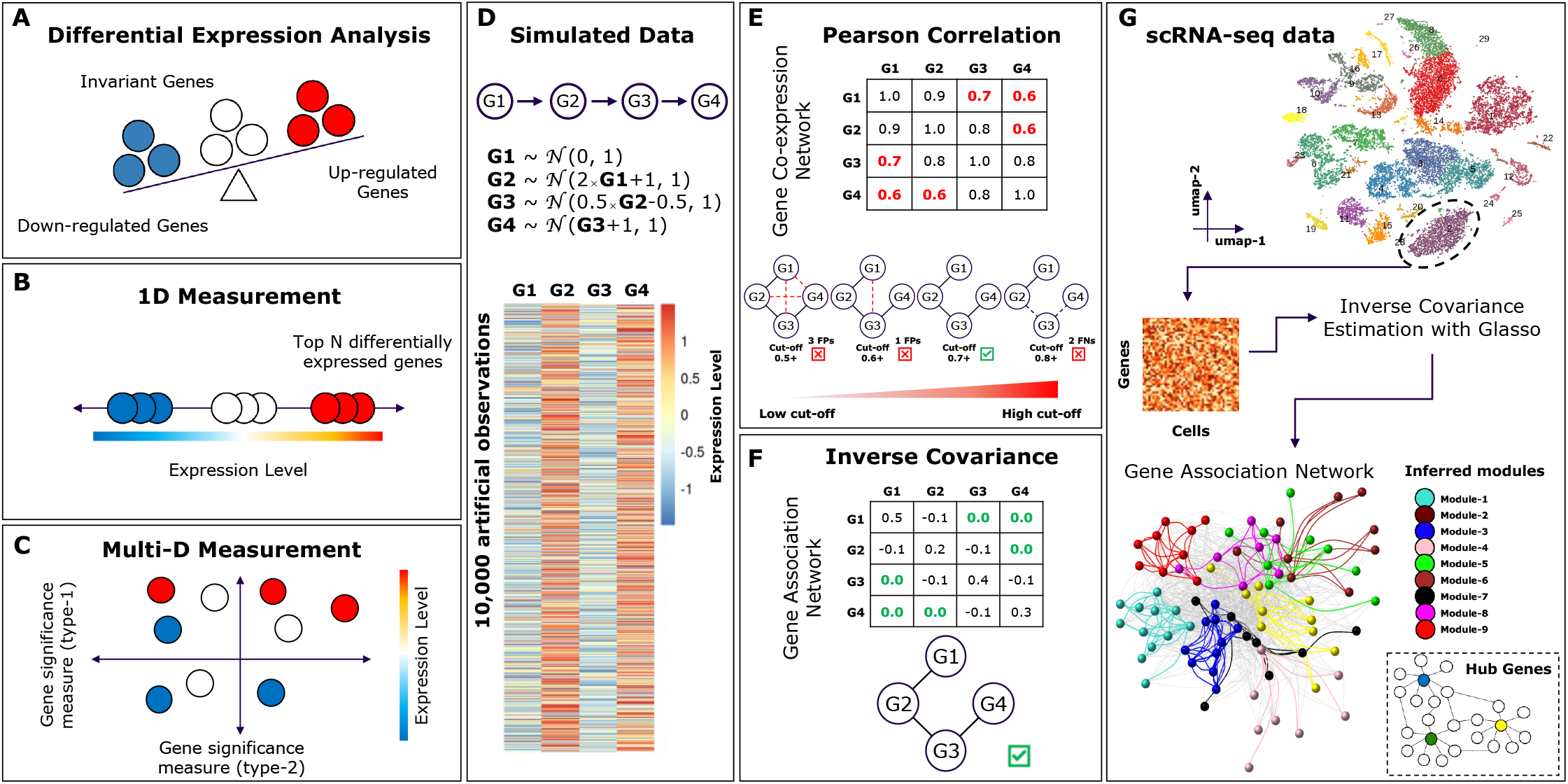
Unveiling overlooked genes through novel measurement techniques. **(A)** Schematic illustration depicting three possible alterations in the expression level of genes upon a stimulus, including upregulation (red circles), downregulation (blue circles), and no variation (white circles). **(B)** Schematic illustration representing a one-dimensional conventional differential expression analysis, aiming to distinguish genes with varying expression levels and identify novel condition-relevant genes. **(C)** Schematic illustration showcasing the potential of novel new measurement techniques to complement traditional differential expression analysis. **(D)** Simulated dataset resembling the sequential regulatory effect of gene G1 on G2, G2 on G3, and G3 on G4. The expression levels are sampled from a Gaussian distribution. The heatmap displays the expression levels of G1-G4 across 10K cells. **(E)** Pearson correlation coefficients calculated from the simulated dataset from **D**, laying the framework to build a gene co-expression network comprised of genes G1-G4. The red elements of the Pearson correlation matrix represent the transitive red edges in the network, i.e., false positives. The networks at the bottom of the panel illustrate the correlation thresholding effort to remove the transitive edges. The red legend at the bottom illustrates the range of thresholds used to remove the transitive edges. **(F)** Inverse covariance matrix is estimated for the same simulated dataset from **D**, laying the groundwork for the inference of a gene association network without any transitive edges, represented by green zeros in the matrix. (**G)** Schematic illustration of the GENIX algorithm starting from single-cell RNA sequencing data. The gene expression matrix of a given cell type is then utilized to estimate the inverse covariance matrix using the glasso algorithm. This process lays the groundwork to build a gene association network and infer associated bio-topological features, hub genes, and modules.

Moreover, there is a growing recognition that the widespread use of statistical significance (often determined by statistical p-value tests) as a basis for making claims of scientific findings may inadvertently mislead the interpretation of results (Amrhein, 2019; Wasserstein, 2019). This concern is particularly salient in the realm of transcriptomics, where genes undergoing substantial changes in expression (determined by log2-fold change) are often classified as significantly differentially expressed if they meet the criterion of an adjusted p-value below a pre-defined cut-off (e.g., 0.05). This issue has recently come under scrutiny in a study that revealed unexpectedly high false discovery rates when identifying differentially expressed genes between two conditions using widely used statistical bioinformatics methods (Li Y. a., 2022). Hence, it is becoming imperative to produce complementary and supporting lines of evidence for scientific inference, going beyond narrow reliance on statistical significance alone.

In this regard, gene networks are informative constructs that can lay the groundwork for a shift towards a multi-layered investigation of the data, where the contextual relevance of genes and their expression changes are given complementary importance. Derived from scRNA-seq data, gene networks provide a robust framework for unraveling the intricate patterns of gene interactions, offering valuable insights into the coordinated behavior and functional relationships within biological systems (Zhang, 2005; Margolin, 2006). Particularly powerful is the application of topological features extracted from these networks to identify essential genes. For instance, measures such as degree, which quantifies the connectivity of a gene within the network, provide indications of its involvement in critical biological pathways. Through such bio-topological measures, we can complement conventional signature identification strategies (**Figure 1C**).

In this study, we introduce GENIX, a novel network-based framework designed to identify genes and gene modules impacted by perturbations across conditions. The approach comprises two main components: (1) a method for sparse, *data-driven* gene association network construction, followed by (2) a measure of individual gene importance with respect to the network topology. In the first component, we apply a probabilistic graphical model to infer undirected dependency graphs, which effectively identifies gene interactions, resulting in biologically meaningful network representations. These networks lay the groundwork for identifying topologically important genes and gene programs. We employ a novel dual metric system for the second component to compare perturbation-driven networks. This systematic approach distinguishes topologically specific genes, whose overall interaction strength is highly condition-specific, from topologically invariant genes, whose connectivity pattern is not predominantly influenced by the perturbation. This novel metric is versatile and can be applied to any sparse network constructed from single-cell data.

In its entirety, GENIX serves as an autonomous ecosystem for network construction. It takes a scRNA-seq count matrix as input to generate undirected dependency graphs alongside a compilation of essential genes and gene modules. Additionally, GENIX provides a unique downstream functionality that enables the comparison of condition-specific inferred networks, effectively allowing for reverse-engineering of perturbations and the identification of novel gene signatures. To demonstrate the power of our approach, we applied it to a previously published time-series single-cell dataset characterizing gender-specific responses to seasonal influenza vaccine in COVID-19-recovered and naïve patients. We reveal the bio-topological properties of the inferred networks in specific cell types and report well-known, as well as novel putative response genes that appear to control differences in protective immunity.

## RESULTS

We evaluated the performance of GENIX in two ways. First, we used synthetic data to demonstrate its ability to faithfully capture built-in relationships in a sparse network. Next, we applied it to a publicly available dataset, GSE206265, reflecting a controlled perturbation (vaccination) in a clinical setting (Sparks, 2023). This dataset consists of longitudinal cellular indexing of transcriptomes and epitopes by sequencing (CITE-seq) data collected at multiple time points (Days 0, 1, and 28) after administrating the seasonal influenza vaccine. It includes samples obtained from individuals who have recovered from COVID-19 (COVR) as well as healthy controls (HC), with a cohort comprising 12 females and 12 males in the COVR group, and 8 females and 8 males in the HC group (**Figure 2A**). Hence, these cohorts encompass 12 groups, representing different genders, sampling time points, and disease conditions. None of the participants were enrolled in COVID-19 vaccine trials and had not received any recent vaccinations prior to receiving the seasonal influenza vaccine.

**Figure 2:**
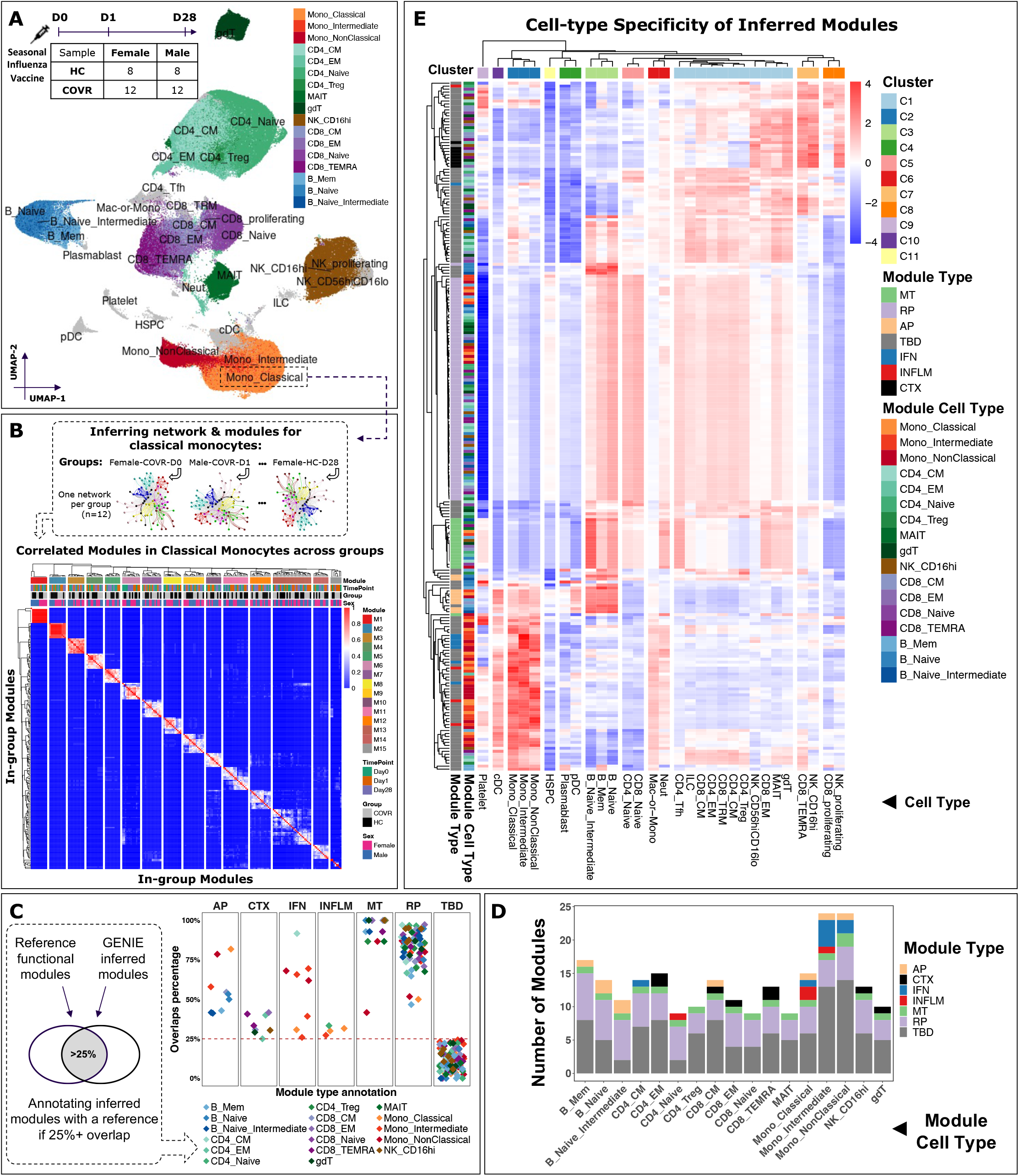
Inferring cell type specific gene modules from query data. **(A)** UMAP representation of GSE206265 PBMC single-cell data obtained from healthy females (n=8), healthy males (n=8), COVID-19 recovered females (n=12), and COVID-19 recovered males (n=12). The groups were vaccinated with the seasonal influenza vaccine, and samples were collected longitudinally at Day 0, 1, and 28 post-vaccination. Clusters and cell types were color-coded according to the published manuscript *(Sparks, 2023)*. **(B)** Schematic illustration of the module inference process for each group of patients (n=12; top panel). The panel showcases only the classical monocyte population. Correlation analysis followed by hierarchical clustering was performed to combine in-group-module gene sets (bottom panel). The color keys on the top represent the clustered modules, sampling time points, disease groups, and genders. Row and column dendrograms are generated by hierarchical clustering. **(C)** Schematic representation for annotating modules by comparing them to the reference functional modules *(Chaussabel, 2008; Chiche, 2014)* (left panel). Modules that have a similarity larger than 25% (red dashed line) with the reference modules are annotated accordingly (right panel). The facet titles on the top represent the modules annotation: AP for antigen presenting, CTX for cytotoxic, IFN for interferon, INFLM for inflammation, MT for mitochondrial, RP for ribosomal protein, and TBD for unannotated modules. The color-coded legend on the bottom represents the cell types from which the modules have been inferred. **(D)** Number of annotated modules (y-axis) in each cell type (x-axis) from which they have been inferred. The legend on the right represents the modules annotation similar to panel **C**. **(E)** Heatmap representing row-scaled module scores (rows) averaged over cells from each cell-type population (columns). The color keys on the left represent the annotated module types and the cell types from which the modules have been inferred. The color key at the top represents the cell type associations with hierarchical clusters.

### GENIX Network Construction Disentangles Direct Associations from Indirect Effects in Synthetic Data

Pearson correlation is a widely adopted method to quantify the relationship between genes and serves as the foundation for constructing networks (Zhang, 2005; Baran, 2019; Iacono, 2019; Rezaie, 2023). However, Pearson correlation quantifies similarity in gene expression patterns without distinguishing between direct and indirect relationships (Marbach, 2012). We underscored this issue with a fundamental yet targeted analysis. We generated a simulated dataset comprising four pseudo-genes, where the expression levels across 10,000 observations were sampled from Gaussian distributions with different characteristics. To emulate a linear relationship among these pseudo-genes, each Gaussian distribution is influenced by the previous one and, in turn, impacts the following one sequentially. The simulation started with a standard normal distribution with a mean of zero and a standard deviation of one (**Figure 1D**). When constructing a gene co-expression network using Pearson correlation, as expected, we observed transitive connections that introduced false positive edges (**Figure 1E**).

One commonly employed approach to address the challenge of transitive connections is correlation thresholding (Zhang, 2005; Care, 2019). However, the choice of the cut-off requires careful consideration, and while it may reduce the impact of transitive connections, it may not completely resolve the bottleneck (**Figure 1E**). Therefore, we opted for an alternative approach by utilizing the inverse covariance matrix (ICM; also known as the precision matrix) (Wasserman, 2004). The ICM offers the advantage of disentangling direct associations between genes from indirect effects mediated through other genes and captures conditional dependencies between variables. In this case, an absent edge means that the two genes are conditionally independent given all other genes and corresponds to a zero element in the ICM. Hence, it reduces the occurrence of false positives, providing a robust and efficient approach for inferring gene networks, referred to as gene association networks (GANs) (**Figure 1F**).

To further validate the relevance of ICM, we extended our simulated pseudo-gene analysis by arranging the pseudo-genes in four distinct configurations, resembling alternative transcriptional relationships. This supplementary analysis highlights the versatile application of the ICM in constructing diverse network structures, effectively addressing the challenge of transitive connections across various contexts (**Supplementary** Fig 1). Therefore, when applied to real single-cell data, ICM captures the partial correlations (see **Methods**) between genes while accounting for the influence of other genes in the network, providing a reliable framework for inferring cell-type-specific gene modules and identifying genes with high centrality (**Figure 1G**).

Estimating the ICM requires the use of graphical least absolute shrinkage and selection operator (glasso) (Friedman, 2008; Meinshausen, 2006) (see **Methods**). This is achieved by maximizing a likelihood function that is subject to sparsity-inducing penalties governed by a regularization parameter. Higher values of this parameter enhance sparsity, whereas lower values diminish it. To determine a justified regularization parameter, we leveraged simulated scRNA-seq data consisting of six samples of 5,000 cells and 10,000 genes each (see **Methods**). We inferred the GAN for each of the simulated datasets across a range of regularization parameters spanning from 0.25 to 2, with an increment of 0.25. Our objective was to discern the impact of different regularization strengths on the resulting number of identified hub genes (see **Methods**). This analysis showed that as the regularization parameter increased, the number of hubs in the inferred gene network progressively decreased (**Supplementary** Fig 2). This observation resonates with the notion that higher regularization values lead to sparser networks by encouraging the identification of fewer significant associations among genes. To ascertain the optimal choice of the regularization parameter, we invoked the elbow-method, which suggests that a turning point in the relationship between the number of hubs and the regularization parameter corresponds to an appropriate level of sparsity while preserving biologically meaningful associations. This analysis led us to select a single regularization parameter (*ρ* = 1) for all stages of our analysis.

### GENIX Infers Heavy-Tailed Networks in Real Single-Cell Data

Incorporating the controlled vaccination dataset (Sparks, 2023), we applied GENIX to infer GANs for each cell type within each group (n=12). Throughout the analysis, we maintained the original clustering and cell type annotation conducted by the authors for consistency (**Figure 2A**). We aimed to infer GANs for cell types that had a minimum of 500 cells within each group, considering this threshold as informative for a reliable analysis, *i.e.*, the given group has enough observations to derive an informative graph (see **Methods** for specific details regarding gene, cell, and group filtration). We specifically inferred GANs for monocytes (classical, intermediate, and non-classical), CD4+ T cells (central memory, effective memory, naïve, regulatory, MAIT, and gamma-delta), CD8+ T cells (central memory, effective memory, naïve, and TEMRA), CD16^high^ NK cells, and B cells (memory, naïve, and intermediate).

Biological networks exhibit non-random characteristics. Specifically, many of these networks demonstrate a distribution of gene degrees that follows a heavy-tailed pattern, with some genes having degrees much higher than the average. Therefore, we sought to investigate this topological property and assure the biological resemblance of the inferred networks. We utilized the networks derived from classical monocytes (n=12) as a showcase. Most genes displayed only a few connections, while a small subset of centralized genes exhibited a disproportionately large number of connections, resulting in a heavy-tailed degree distribution (**Supplementary** Fig 3). This analysis underscores the biological authenticity of the inferred networks, aligning them with patterns commonly observed in intricate biological systems (Stumpf, 2012; Holme, 2019).

### GENIX Infers Modules Related to Biological Function Within and Across Cell Types in Real Single-Cell Data

Modularity is a key property of biological networks. To explore this, we examined graphs inferred from classical monocytes in each patient group (n=12) and identified in-group modules (i.e., sets of densely interconnected genes; see **Methods** and **Figure 2B**). Next, we sought to construct unified cell type-specific modules from the in-group modules. We calculated the Jaccard Index to measure the level of similarity among the identified in-group modules. We then applied hierarchical clustering to identify clusters of modules with similar genes (**Figure 2B**). Henceforth, the term ’module’ signifies these resultant clusters, which represent the union of clustered in-group-modules. We repeated this process for all 17 investigated cell types, resulting in the average identification of 13 ± 4 (SD) modules per each cell type, with a total of 232 modules (**Supplementary File 1**).

Genes within the same module often participate in common biological functions. Therefore, we next sought to elucidate the biological function of each inferred module. We utilized a well-established module set (Chaussabel, 2008; Chiche, 2014) and annotated the inferred modules with the reference functional modules if they exhibited an overlap of above 25% (**Figure 2C**). More specifically, we focused our analysis on three main biological functional modules from the reference dataset: interferon-related modules (M1.2, M3.4, and M5.12), inflammatory-related modules (M4.2, M5.1, M6.13), and cytotoxic-related modules (M3.6, M4.15, and M8.46), which we annotated as IFN, INFLM, and CTX, respectively. Additionally, we annotated inferred modules with above 25% overlap with mitochondrial-related genes, antigen-presenting-related genes (HLAs), and ribosomal proteins as MT, AP, and RP, respectively. Those remaining are annotated as TBDs (**Supplementary File 2**). Finally, we conducted gene enrichment analysis (see **Methods**) as a complementary analysis to annotate module function, leveraging gene ontology (GO) terms (**Supplementary File 3**).

Intriguingly, the modular analysis revealed a cell type-specific attribution of biological functions to the inferred modules (**Figure 2D**). AP modules were predominantly found in antigen-presenting cells, including B and monocyte subtypes. CTX modules were present in diverse killer cells, such as CD4 effective memory cells, CD8 central and effective memory cells, Terminally Differentiated Effector Memory CD8+ T cells (TEMRA), CD16^high^ natural killer cells, and gamma-delta T cells (gdT). IFN and INFLM modules were prominently present in monocyte subtypes. RP and MT modules were identified across all cell types, indicating their fundamental roles in various cellular processes.

Finally, we aimed to demonstrate the module’s specificity concerning all cell types in the dataset. Using the identified modules, we assigned scores to individual cells based on the expression levels of the associated module gene lists (see **Methods**). This analysis allowed us to assess the relative significance of modules within each cell type, categorizing them into 11 groups sharing similar phenotypic characteristics through hierarchical clustering (**Figure 2E**). The largest group, C1 (n = 12), comprised T and NK cell states. C2-3 (n = 3, each) consisted of monocyte and B cell states, respectively. C4-8 (n = 2, each) encompassed plasmatic, naïve CD4 and CD8, neutrophils and macrophages, activated CD8 and NK, and proliferating CD8 and NK states, respectively. Finally, C9-11 (n = 1, each) included platelets, conventional dendritic cells (cDCs), and hematopoietic stem and progenitor cells (HSPCs), respectively. This investigation confirmed the phenotypic specificity of the identified modules. Overall, this comprehensive analysis provided evidence of the biological plausibility of the inferred graphs and the associated identified modules, thus affirming the reliability of GENIX performance.

### GENIX Module Analysis Across Conditions Provides Insights into Variation in Response to Influenza Vaccine

One of the main objectives in time-series vaccination studies is to identify sets of functionally coherent genes and their associations with the immune states (pre-and post-vaccination), thereby providing insights into the essential pathways governing the immune response. Notably, responses to influenza vaccination have been extensively characterized, revealing the augmented activity of the IFN, CTX, and AP pathways post-vaccination (Wimmers, 2021; Mellett, 2022; Giacomelli Cao, 2022). Therefore, vaccination is an ideal perturbation for validating the functional relevance of modules inferred by GENIX.

The longitudinal influenza vaccination conducted by Sparks et al. (Sparks, 2023) reported that male individuals who had recovered from COVID-19 exhibited higher early, influenza vaccine-induced responses compared to healthy male and female individuals, as well as female individuals who had recovered from COVID-19 (COVR). In this context, our investigation aimed to assess the condition specificity of inferred modules among female (F) and male (M) individuals in both the COVR and HC groups. Additionally, we sought to parse out the differences in response in terms of specific driver genes and pathways, thereby providing supporting evidence and enhanced insights into gender-specific responses to immunization.

Given that vaccination induces an antiviral-like response involving both the myeloid and lymphoid compartments, we narrowed our focus to classical monocytes (cMon) and CD8+ central memory T cells (CD8-CM), as these cell types play pivotal roles in orchestrating the early immune response. Notably, the gene enrichment analysis of the inferred modules from cMon and CD8-CM populations (as discussed in the previous section) recapitulated essential immunoregulatory functions (**Supplementary File 2**). Specifically, the cMon.M2 module exhibited enrichment of AP-related genes, including major histocompatibility complex (HLA) class-II genes, while the cMon.M15 module demonstrated enrichment with IFN-related genes. In addition, the CD8-CM.M7 module showed enrichment of CTX-related genes, and CD8-CM.M9 displayed enrichment of AP-related genes, including HLA class-I genes (**Supplementary** Fig 4). We leveraged these immune-response driving modules to score each cell (see **Methods**) and explore their comparative importance among cMon and CD8-CM populations. This analysis highlighted the cell-type specificity of the inferred IFN and CTX modules, effectively distinguishing cMon and CD8-CM cells (**Figure 3A** - top panel), underscoring their unique functionality in modulating immune responses. Additionally, the comparison between cMon and CD8-CM cells, scored using HLA class-I and II modules, emphasized the phenotypic association of the HLA class-II molecules to antigen-presenting cells (**Figure 3A** - bottom panel).

**Figure 3:**
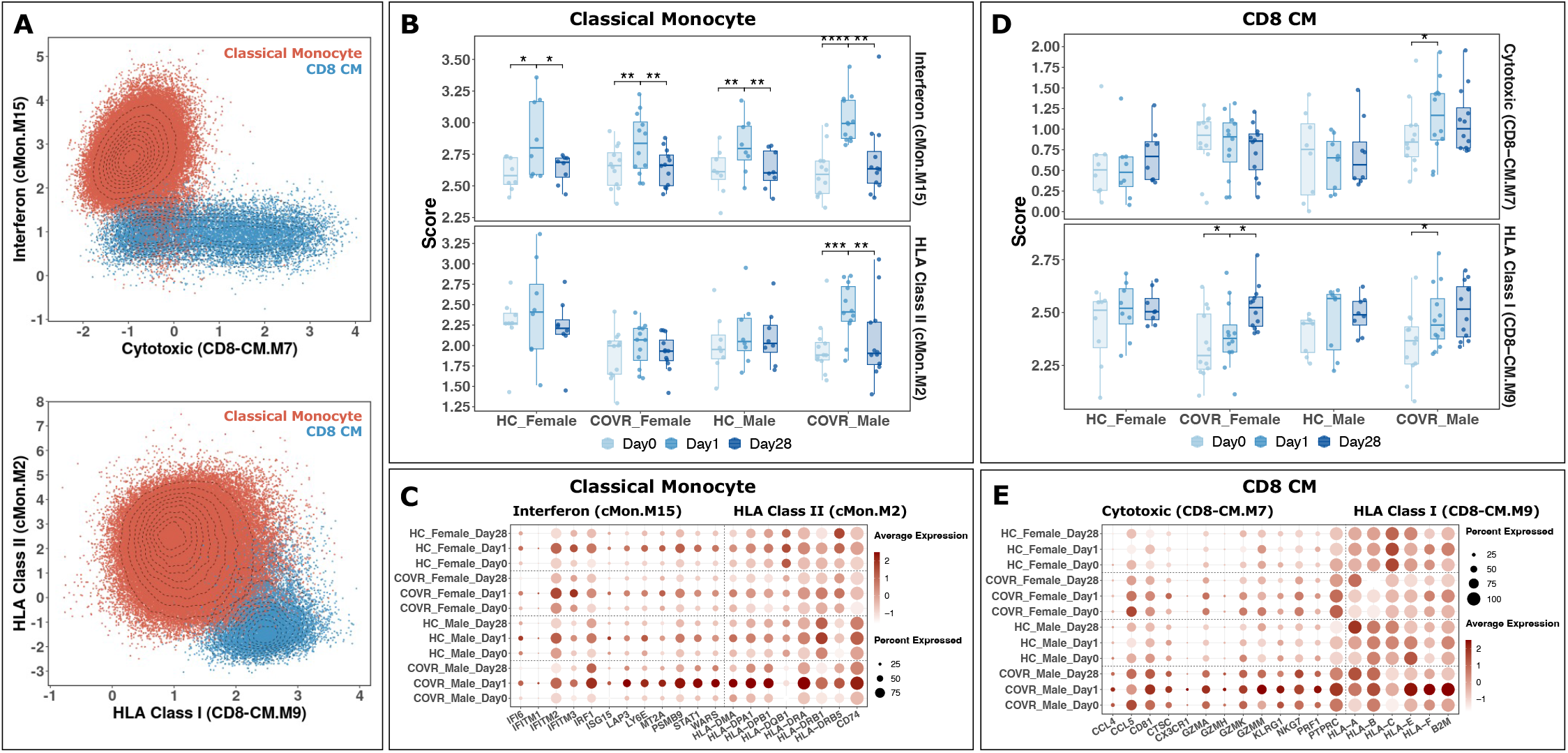
Discovery of vaccine-induced gene programs in classical monocytes and CD8 central memory cells. **(A)** Visualization of classical monocytes (dark orange) and CD8 central memory cells (dark blue) in the context of the inferred interferon (cMon.M15) and cytotoxic (CD8-CM.M7) modules in the top panel, and HLA class II (cMon.M2) and HLA class I (CD8-CM.M9) modules in the bottom panel. **(B)** Module scores of interferon (cMon.M15) and HLA class II (cMon.M2) modules in classical monocytes in the HC-Female (n = 8), HC-Male (n = 8), COVR-Female (n = 12), and COVR-Male (n = 12) participants at D0, D1, and D28. Each point represents a participant. The upper and lower bounds of the boxplot represent the 75th and 25th percentiles, respectively. The center bars indicate the medians, and the whiskers denote values up to 1.5 interquartile ranges above the 75th or below the 25th percentiles. Data beyond the end of the whiskers are considered outliers. **(C)** Expression levels of selected genes from the interferon (cMon.M15) and HLA class II (cMon.M2) modules in classical monocytes across different groups and time points. Each circle’s color represents the average expression level of the gene, while its size indicates the percentage of cells expressing that particular gene. **(D-E)** Similar to panels **B** and **C**, but calculated from CD8 central memory cells using cytotoxic (CD8-CM.M7) and HLA class I (CD8-CM.M9) modules.

We hypothesized that the activity of these modules may vary in a sex-and time-dependent manner in the context of vaccination and that this variation may explain differences in response across groups. We therefore investigated the time-dependent score of these modules in each group. At baseline, pre-vaccination IFN-related activity (cMon.M15) demonstrated consistency among all participants (**Figure 3B**). Additionally, a significant increase in IFN activity was observed in cMon on Day 1 after influenza vaccination. The elevated IFN activity returned to normal at Day 28 across all groups. Notably, the increase in IFN activity captured by cMon.M15 was particularly pronounced in the COVR-M group. Concurrently, a significant increase in the activity of the MHC class II module (cMon.M2) was observed only in the COVR-M group on Day 1 (**Figure 3B**). Thus, our module-based analysis highlights an amplified response in the COVR-M group, compared to other patient groups, evident within the cMon population on Day 1 following seasonal influenza vaccination. This heightened activity is notably concentrated in the IFN module, encompassing genes coding for IFN-inducible antiviral proteins such as *IRF1*, *ISG15*, and *IFITM3*, as well as in the enhanced AP activity, characterized by elevated expression of HLA class-II related genes including *HLA-DRA*, *HLA-DMA*, and *CD74* (**Figure 3C**).

Similarly, a significant increase in the activity of the CD8-specific CTX module (CD8-CM.M7) was observed in COVR-M individuals on Day 1 after influenza vaccination (**Figure 3D**). This elevated activity was suppressed by Day 28. Additionally, a relatively modest response in the activity of the MHC class I module (CD8-CM.M9) was observed on Day 1 across all groups (**Figure 3D**), with the exception of the COVR groups, which exhibited a particularly pronounced increase again in male patients. Thus, the COVR-M group exhibits an elevated CTX response in CD8-CM T-cells, characterized by increased expression of key cytotoxic genes such as *PRF1*, *GZMK*, *GZMM*, *KLRG1*, and *NKG7* (**Figure 3E**), accompanied by an enhanced AP activity, marked by elevated expression of HLA class-I related genes such as *HLA-A*, *HLA-B*, *HLA-C*, and *B2M* (**Figure 3E**), on Day 1 following seasonal influenza vaccination.

In summary, these results are consistent with the findings in the original paper (Sparks, 2023), indicating that male individuals who had recovered from COVID-19 exhibited coordinately higher early, influenza vaccine-induced responses compared to other groups in the study. In addition, GENIX identified the IFN, CTX, and AP cell-type-specific modules as drivers of this difference, thereby providing supportive and detailed insights into the underlying immune pathways at play following seasonal influenza vaccination.

### GENIX Identifies Vaccine-Response Gene Signatures Beyond Differential Expression

To identify genes that drive group-specific vaccine responses, we devised a two-dimensional topological measure that compares the structure of pre-and post-vaccination inferred networks. Prior to this, we sought to establish that the observed differences in network structures are not mere artifacts of random noise, but rather indicative of significant variations. To achieve this goal, we employed a permutation test (see **Methods**). This test aimed to gauge the robust distinctiveness of inferred networks from cMon at D0 and D1 after vaccination in individuals who had recovered from COVID-19. By juxtaposing the observed Jaccard similarity index between pre-and post-vaccination networks with corresponding indices calculated from randomly reconstructed networks (50,000 times), our analysis underscored the statistical significance (p < 0.05) of structural disparities between the observed networks (**Supplementary** Fig 5). This substantiated the identification of reliable topological changes stemming from perturbations.

To measure the importance of a given gene in the overall structure of the network, we formulated a unique metric that gauges the consequences of gene knockout on the network topology pre-and post-vaccination (see **Methods**). We started by computing the similarity of networks across conditions using their Jaccard index, considering both shared genes and edges (**Figure 4A**, panel 1). Next, we conducted an iterative removal of each gene from both networks and recalculated the similarity index. Finally, we defined the gene removal impact (GRI) metric of each gene as the relative difference in similarity of networks after removal compared to before removal. The GRI metric can take both positive and negative values, underscoring its potential to discern genes substantially impacting on the network structure. A positive GRI indicates that the elimination of a specific gene leads to a more similar state between the networks (**Figure 4A**, panel 2), while a negative GRI indicates that the elimination of a specific gene causes the networks to become less similar (**Figure 4A**, panel 3).

**Figure 4:**
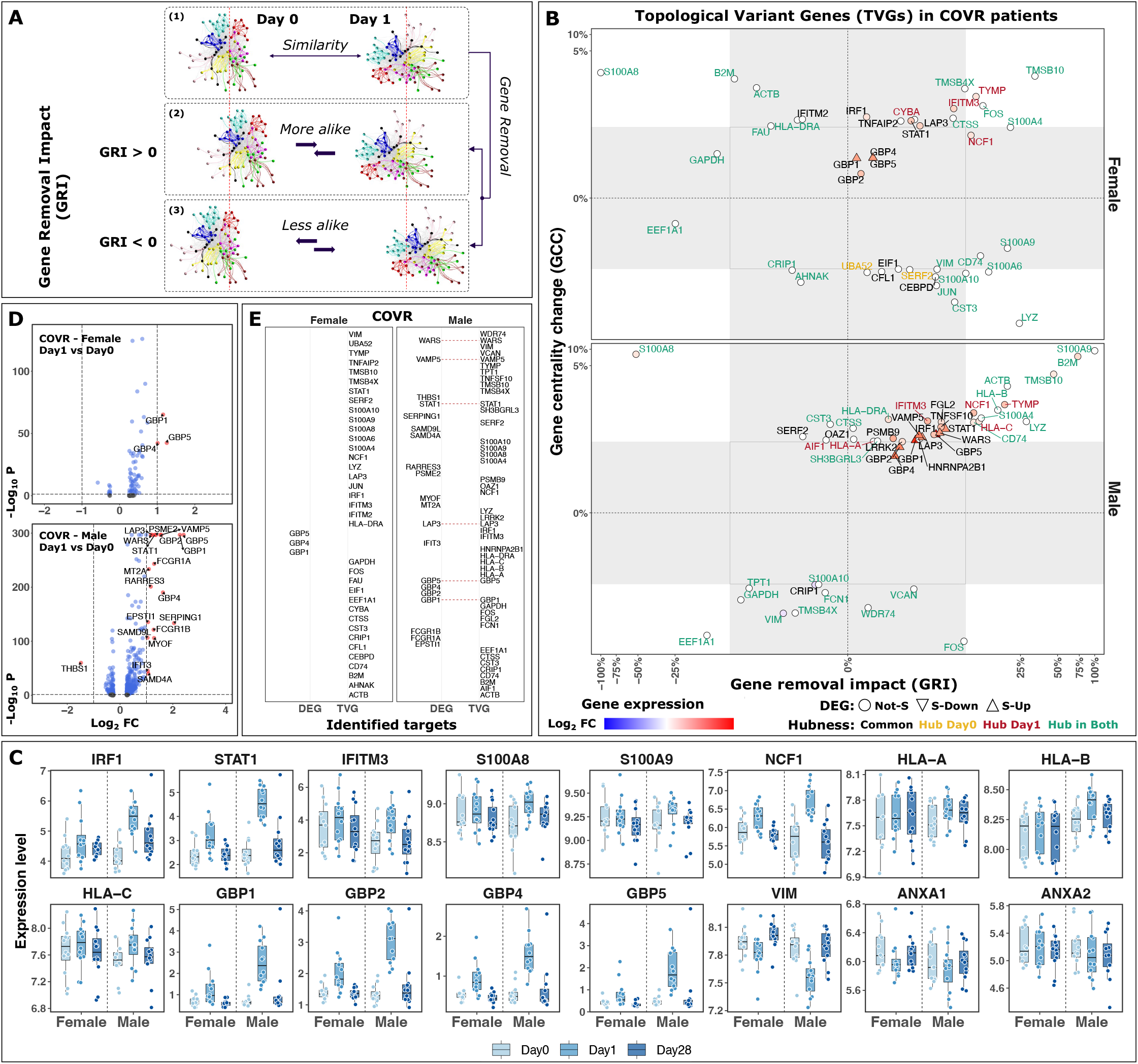
Identification of vaccine-induced topological variant genes (TVGs) in classical monocytes. **(A)** Schematic illustration of the gene removal impact (GRI) metric calculation. The Jaccard Index is calculated to quantify the similarity, considering both shared genes and edges, comparing the inferred networks at D0 and D1 post-vaccination (panel 1). Genes are iteratively removed from both networks, and the similarity index is recalculated. The impact of individual genes is then determined through the relative comparison of these pre-and post-removal measurements. The GRI is positive if the elimination of a specific gene leads to a more similar state between the two networks (panel 2). The GRI is negative if the elimination of a specific gene causes the two networks to become less similar (panel 3). **(B)** Change in gene normalized centrality (y-axis; D1 versus D0) is compared against gene removal impact (x-axis; D1 versus D0) in inferred networks from classical monocytes in female (top panel) and male (bottom panel) participants recovered from COVID-19 (COVR). Each point indicates a TVG labeled by the gene name. The label color indicates the hubness status of the given gene, either being a hub at D0 (orange), D1 (red), in both D0 and D1 (green), or not a hub gene (a common gene) in neither time points (black). The points color shade indicates the log-2-fold change (Log2 FC) in the expression level of the given gene, comparing D1 post-vaccination versus D0. The point shape indicates the differential expression status of each individual gene, either being invariant (circle), significantly downregulated (inverted triangle), or significantly upregulated (triangle). The shaded area indicates the region where the removal of the given gene has a low influence on the overall architecture of the inferred networks, quantified with a change less than 3-sigma in either the GRI (x-axis) or GCC (y-axis) distributions. The x-axis values are normalized to their extremum positive and negative values. **(C)** Expression levels of selected genes in classical monocytes of COVID-19 recovered female (n = 12) and male (n = 12) participants at D0, D1, and D28. Each point represents a participant. The upper and lower bounds of the boxplot represent the 75th and 25th percentiles, respectively. The center bars indicate the medians, and the whiskers denote values up to 1.5 interquartile ranges above the 75th or below the 25th percentiles. Data beyond the end of the whiskers are considered outliers. **(D)** Volcano plot illustrating differential expression analysis (DEA) comparing D1 post-vaccination versus D0 in classical monocytes from female participants (top panel) and male participants (bottom panel) recovered from COVID-19 (COVR). Red points represent genes with statistically significant up-or down-regulation (p-adjust less than 0.05 and absolute log2 fold change larger than 1), while blue points correspond to genes satisfying only the statistically significant criteria (p-adjust less than 0.05). Black points denote genes that do not meet either of the two criteria. **(E)** Shows vaccine-induced gene signatures identified using both the DEA and the network-based comparative model in female and male participants recovered from COVID-19 (COVR). The dashed red line indicates signatures identified by both methodologies as differentially expressed genes (DEGs) or topological variant genes (TVGs). Note: the GBP1,2,4, and 5 genes have not been removed from the panel **B** to highlight the complementary nature of DEA and network-based comparative model in identifying perturbation-induced signatures.

To complement the GRI metric, we also incorporated the change in gene centrality to enhance our analysis. By comparing the normalized degree of connectivity for each gene within the pre-and post-vaccination inferred networks, we derived the gene centrality change (GCC) metric (see **Methods**). In this context, a positive GCC indicates that specific genes become more central to the network structure, whereas a negative GCC suggests that certain genes become less central, signifying a potential shift in their importance or regulatory influence following vaccination.

Concentrating our analysis on classical monocytes from individuals who had recovered from COVID-19, we identified 36 topologically variant genes (TVGs) (see **Methods**) in the female group and 44 TVGs in the male group, when comparing inferred networks from Day 1 post-vaccination to Day 0 (**Figure 4B**). Several interferon-related genes, including *IRF1*, *STAT1*, and *IFITM3* (part of interferon module cMon.M15) were identified in both groups (**Figure 4B-C** and **Supplementary File 1**). Notably, the type I interferon-inducible protein 3 (*IFITM3*) emerged as a hub gene solely at Day 1 after influenza vaccination, suggesting its potential role in the early response to vaccination (Cao, 2014).

Furthermore, we detected inflammatory-related genes such as calprotectin *S100A8/9* (belonging to inflammatory module cMon.M4) and *NCF1* (belonging to inflammatory module cMon.M14) in both male and female groups (**Figure 4B-C** and **Supplementary File 1**). Noteworthy is the neutrophil cytosolic factor 1 (*NCF1*), a regulator of reactive oxygen species (ROS), which exhibited hub gene characteristics exclusively on Day 1 post-vaccination, emphasizing its potential immunoregulatory effects in response to the seasonal influenza vaccine. Additionally, the HLA class-I gene set (A, B, and C), all belonging to module cMon.M7 and demonstrating hub gene characteristics on Day 1 post-vaccination, were uniquely observed in the male group (**Figure 4B-C** and **Supplementary File 1**).

Next, we sought to juxtapose the outcomes of our network-based comparative model with those of the DEA in uncovering vaccine-induced signatures. We carried out the DEA using time point D0 as the baseline in classical monocytes from COVID-19 recovered female and male participants. We stablished a threshold of adjusted p-values below 0.05 and a log-2-fold change exceeding 1. This led us to identify three genes in the COVR-F group and 20 genes in the COVR-M with significantly altered expression patterns at Day 1 post-seasonal influenza vaccination (**Figure 4D**).

Interestingly, none of the three differentially expressed genes, *GBP1*, *GBP4*, and *GBP5* from the COVR-F group were detected by the network-based comparative model (**Figure 4E**). In fact, all three of them resided within the non-essential region **(Figure 4B** and **4C**). Conversely, all 36 genes identified by the network-based comparative model were overlooked by the DEA, despite comprising several key immunoregulator genes known to mediate response to vaccination and various cellular stimuli, including interferon-related genes *IRF1*, *IFITM2*, and *IFITM3* (Feeley, 2011; Zhai, 2015; Keshavarz, 2019; Wimmers, 2021), as well as inflammatory-related genes *S100A8*, *S100A9*, and *NCF1* (Peiris, 2009; Tsai, 2014; Holmdahl, 2016; Mellett, 2022).

Moreover, in the COVR-M group analysis, both methodologies converged on 6 common genes, 5 of which were interferon-related, including *WARS*, *STAT1*, *LAP3*, *GBP1*, and *GBP5* (**Figure 4E**). Interestingly, the immune-inflammatory regulator vimentin (*VIM*) (Li, 2020), absent from the DEA, was unveiled through network analysis in both the female and male groups (**Figure 4E**). Notably, *VIM* was also a constituent of the inferred cMon.M3 module, which included Annexins A1 and A2 (*ANXA1/2*) recognized for their role in influenza virus pathogenesis (Ampomah, 2018; Rahman, 2018) (**Supplementary File 1** and **Supplementary** Fig 4A). This observation underscores the potential immune regulatory coherence of *VIM* and *ANXA1/2*, rendering them intriguing candidates for further investigation, especially within the context of influenza vaccination (**Figure 4C)**. Overall, this comparative analysis highlights the complementary nature of both methodologies in unveiling perturbation-induced signatures.

## DISCUSSION

In the pursuit of unraveling the complexities of biological systems, traditional identification of gene regulatory responses approaches, such as DEA, have proven to be indispensable tools. However, these approaches are primarily designed to prioritize genes that exhibit significant changes in their expression levels, potentially overlooking other genes that may play crucial roles in the studied biology despite not showing pronounced alterations in expression. To address this limitation and enhance our comprehension of the intricate interplay within biological systems, we have developed a novel network-based comparative model, integrated into GENIX algorithm. This approach focuses on condition-driven structural changes within gene networks, thereby highlighting genes that might have otherwise gone unnoticed through conventional analyses but could be essential components of the underlying biological pathways driving a specific response to perturbation.

Specifically, we utilized a glasso-based network construction approach to capture gene expression dependencies in single-cell data. By leveraging this probabilistic graphical model, GENIX faithfully differentiates between direct and indirect connections while remaining immune to neglecting novel interactions, a common downside of reference-guided network construction methods (Mohammadi, 2019). Within GENIX, we further developed a systematic module identification and analysis approach, and a two-dimensional quantitative metric, providing a more comprehensive understanding of changes in gene essentiality within the network upon perturbation. We introduced the gene removal impact (GRI) metric, which assesses the influence of a specific gene on the overall change in architecture of inferred networks across conditions. The GRI is combined with the gene centrality change (GCC) metric, which measures the degree of a specific gene’s involvement in response to the perturbation. These metrics can yield positive and negative values, highlighting their potential to distinguish topologically variant genes (TVGs) that impact the network’s structure.

To provide biological intuition to the GRI metric, we envision a 3D space wherein the pre-perturbation network state is located at the center, while the post-perturbation network state resides at a distinct location in space, symbolizing the alterations brought about by the perturbation. In this context, a positive GRI indicates that removing a specific gene leads to a convergence of the networks by increasing their structural similarity. This convergence suggests that the biological pathway involving the gene is likely to drive the changes associated with the perturbation and that knocking out the gene would reverse or attenuate the effect of the perturbation. Inversely, a negative GRI indicates that removing a specific gene causes the networks to diverge. This divergence suggests that the relationship between the biological pathway involving the gene and the perturbation is likely complex and multifaceted and that knocking out the gene would amplify the effect of the perturbation.

With regard to the GCC metric, a positive change in centrality implies the establishment of new connections or interactions within the network, serving as a proxy for revealing novel regulatory relationships that emerge in response to the perturbation. Conversely, when GCC is negative, the reasoning is the opposite, suggesting that certain genes may interact less or become more specific in the altered pathways. The GCC and GRI metrics are orthogonal approaches to assess the importance of individual genes, and by integrating them, we have provided a comprehensive view of how individual genes influence the overall network following a biological perturbation.

Conducted within the context of the seasonal influenza vaccine, we assessed the applicability and performance of GENIX. We inferred cell-type-specific and condition-specific gene association networks from healthy individuals and COVID-19 recovered individuals who underwent vaccine administration. Subsequently, we analyzed the bio-topological properties, including modules and hubs, of the inferred networks. Our findings demonstrated the cell-type specificity of gene modules and their biological functional coherency, supporting their immunobiological plausibility. Furthermore, we reported both well-known and novel regulatory signatures that might have implications for protective immunity against influenza. Finally, we demonstrated the effectiveness of our novel network-based comparative model in complementing traditional signature identification approaches, such as differential expression analysis.

The glasso algorithm employs a tuning parameter (*ρ*) to modulate the sparsity level of the inferred network. A common approach to selecting the glasso tuning parameter involves constructing multiple networks across a range of *ρ* values and identifying the most suitable value based on its impact on a network topological criterion (Zhao, 2006; Epskamp, 2018). We used the count of identified hubs as a surrogate to assess the impact of different *ρ* values, ultimately selecting a value that exhibited a turning point in the changes of the total hub count as larger values were considered. An alternative approach for tuning parameter determination involves minimizing the extended Bayesian information criterion (EBIC) (Chen, 2008). Despite EBIC’s generally high specificity, its sensitivity can display variability (Foygel, 2010; Van Borkulo, 2014; Barber, 2015). Not surprisingly, EBIC is once again dependent on another hyperparameter, which governs the extent of preference for simpler models within EBIC’s framework. Thus, determining the optimal hyperparameters is tied to the characteristics of the data under investigation. Effective selection of the glasso tuning parameter, tailored to the unique attributes of scRNA-seq data, will remain an important area of further study.

The glasso algorithm employs a streamlined approach to enhance computational efficiency (Witten, 2011). In terms of the computational load, utilizing a single core with a 3.1 GHz processor and 128 GB memory, we recorded that the process of constructing a network with 10,000 genes expressed across 5,000 cells (the size of simulated data used in this study) may take around 5 hours. However, it is worth noting that a pre-filtering stage can help improve the run-time. In particular, a considerable number of genes are expressed sparsely across cells, and these genes can be removed as they do not provide sufficient observations to participate in the network inference. Another approach is to focus only on genes with high variations across conditions.

However, this method may eliminate genes with robust expression across conditions, concentrating solely on genes drastically impacted by perturbation, potentially defeating the purpose of our network-based approach.

It is noteworthy that mutual information (MI) has also been proposed as a basis for network construction (Margolin, 2006). However, while MI is well-defined for discrete or categorical variables, estimating it between quantitative variables poses challenges and can be computationally intensive. In addition, independent studies have shown that MI often falls short compared to correlation-based methods in revealing gene pairwise relationships and identifying co-expression modules (Song, 2012). Each network inference method is underpinned by specific theories and assumptions that may introduce biases in estimating network topologies. Thus, conducting a comprehensive and independent evaluation of these methodologies necessitates additional scrutiny.

Deep learning architectures have been developed for predicting transcriptional responses to single-gene perturbations (Lotfollahi, 2019; Roohani, 2023). However, their predictive accuracy is constrained by the need for reliable training data and prior knowledge of gene-gene relationships. GENIX can serve as a valuable complement, enriching these emerging *in silico* technologies by supplying essential information, including data-driven gene-gene graphs, as well as biologically plausible target genes and gene programs. This alliance enhances the capabilities of predictive modeling, aimed at achieving a spectrum of outcomes with greater precision.

In conclusion, our integration of insights from gene networks and advanced quantitative metrics has provided a comprehensive perspective on how individual genes exert influence over the network structure following perturbation. Our approach complements traditional signature prioritization methods and deepens our understanding of the underlying biological processes. Hence, in conjunction with other lines of evidence, GENIX has the potential to inform target identification and response biomarker discovery pipelines by pinpointing genes with pivotal roles in perturbation-induced pathways from scRNA-seq data.

## ACKNOWLEDGMENT

We would like to express our gratitude to M. Todd Valerius, Franck Rapaport, and Andre H. Kurlovs from Sanofi Precision Medicine and Computational Biology, as well as to Ziv Bar-Joseph and Sachin Mathur from Sanofi R&D Data & Computational Sciences for their valuable comments on the manuscript. We would like to acknowledge Matthew Chamberlain, a former member of the group, for his initial investigations of single-cell network construction using the glasso methodology. This work was supported by Sanofi US.

## AUTHOR CONTRIBUTION

**NN, GG**, **HM**, and **VS**: Conceptualization. **NN** and **VS**: Writing Original Draft of the Manuscript. **NN**: Design, Methodology, Software, and Analysis. **NN**, **GG**, **HM,** and **VS:** Reviewing, Editing, and Extending the Manuscript. **VS** and **EdR**: Supervision and Strategic Direction.

## DEDCLARATION OF INTEREST

The authors are employees of Sanofi US.

## SUPPLEMENTAL INFORMATION

Supplemental information can be found online at Cell Systems.

## DATA AND CODE AVAILABILITY

The GENIX R package source code has been uploaded to GitHub at https://github.com/Sanofi-Public/PMCB-Genix and is publicly available. This paper analyzes existing, publicly available data. Single-cell data are available from the NCBI Gene Expression Omnibus, accession numbers GSE206265. The source code and materials required to reproduce the results will be publicly available as of the date of publication.

## METHODS

### Cells and Genes Pruning

Given the vast number of genes measured in scRNA-seq experiments, it is prudent to apply filtering strategies to focus on a more informative subset of genes and cells. In our study, prior to inferring networks, we applied filters to exclude genes expressed in less than 10 percent of the cells and cells expressing fewer than 250 genes. Cell populations with less than 500 cells per given condition were also excluded from the analyses. Accordingly, we excluded CD4_Tfh, CD8_TRM, CD8_proliferating, NK_proliferating, NK_CD56hiCD16lo, Plasmablast, Mac_or_Mono, Neut, HSPC, pDC, cDC, ILC, and Platelet populations. Additionally, we removed from the analyses small doublet populations, such as CD4_platelet_bind and Mono-T-dblt.

### Count Matrix Normalization

The count expression matrix is normalized to the mean library size, where each cell is scaled to sum up to the mean total counts.

### Inverse Covariance Matrix (ICM) Estimation

The glasso algorithm was utilized to estimate the ICM (Friedman, 2008; Meinshausen, 2006). This method uses a regularization technique that extends the concept of ordinary least squares regression to estimate the ICM, of which the elements are proportional to the partial correlations between variables (Yuan, 2007; Banerjee, 2008; Mazumder, 2012). The “glasso” function from the glasso R package (version 1.11) was used to estimate the inverse covariance matrix (Friedman, 2008) with a regularization parameter value of 1. The default arguments were employed for all the analyses in this study.

### Partial Correlation Matrix Calculation

When considering the relationship between the graphical lasso and partial correlation, let *R* denote the matrix of partial correlations and *Ώ* represent the corresponding inverse covariance matrix, then the *i*^*th*^, *j*^*th*^element of *R* can be computed as *R*_#,%_ = −*Ώ*_#,%_⁄)*Ώ*_#,#_*Ώ*_%,%_.

### scRNA-seq Data Simulation

The Splatter R package (version 1.24.0) (Zappia, 2017) was used to create a set of 6 simulated scRNA-seq data with 5,000 cells and 10,000 genes. The default arguments were employed for all the analyses in this study.

### Module Inference

First, each network is represented as an adjacency matrix A. To normalize the data, the values are shifted from [−1, 1] to [0, 1] using the formula *A*^1^ = 0.5 + 0.5 × *A*. Subsequently, the transformed adjacency matrix is converted into a distance measure by subtracting each value from 1, (1 − *A*^1^). Next, the distance matrix is subjected to hierarchical clustering using the “average” method. This clustering approach generates a dendrogram, allowing for the identification of cohesive modules. Finally, the modules are automatically delineated based on the dendrogram’s structure using the “cutreeDynamic” function from the dynamicTreeCut R package (version 1.63.1), with a minimum module size of 7. The default arguments were employed for all the analyses in this study.

### Genes Centrality Calculation

The degree of a gene in the network is calculated as the number of direct connections the gene has with other genes. A normalized degree for the gene is calculated by dividing the gene’s degree by n-1, where n is the number of vertices in the network.

### Hub Identification

A gene is classified as a hub if its degree value is an outlier, exceeding at least 1.5 interquartile ranges above the 75th percentile of all degrees in the network and if it is connected to at least 1% of the total genes in the network.

### Networks Distinctiveness

A permutation test is developed to examine the statistical distinctiveness of the overall structures of the inferred networks before and after perturbation. The process is initiated by calculating the Jaccard index to assess network similarity, taking into account shared genes and edges. This resulting value serves as the observed statistic. Subsequently, the two networks are combined (union) into a pull, providing the foundation for subsequent comparisons. Pre-and post-perturbation networks are then reconstructed by randomly sampling from the pull, while preserving the original edge counts and connections, with no-replacement. Through an iterative process of resampling and calculating the Jaccard index, a distribution of similarities is generated (n=50,000). This distribution is utilized to calculate a two-sided t-test p-value, reflecting the proportion of permuted statistics that differ from the observed statistic. A p-value less than 0.05 is employed to classify the statistical significance of structural disparities between gene networks before and after perturbation.

### Gene Removal Impact (GRI)

Pre-and post-perturbation network similarity is calculated using the Jaccard index, considering both shared genes and edges. An iterative gene removal process is then executed in both network states. With each gene removal, the similarity index between network states is recalculated. The GRI for each gene is calculated as the relative difference in post-removal network similarity compared to the pre-removal state.

### Gene Centrality Change (GCC)

The GCC for each gene is calculated by subtracting the normalized degree of genes between pre-and post-perturbation network states. A normalized degree for the gene is calculated by dividing the gene’s degree by n-1, where n is the number of genes in the network. Genes that were not included in at least one of the networks are assigned a degree value of zero.

### Topologically Variant Genes (TVG)

A gene is classified as topologically variant if it exhibits a change in either the GRI or GCC metrics exceeding three times the standard deviation (3-sigma) of the distribution of GRIs and GCCs.

### Single-cell Module Score Calculation

The “AddModuleScore” function from Seurat R package (version 4.3.0) was used to calculate module scores for each individual cell based on the average expression of associated gene lists. A positive score would suggest that this module of genes is expressed in a particular cell more highly than would be expected, given the average expression of this module across the population. The default arguments were employed for all the analyses in this study.

### Gene Enrichment Analysis

Enrichment between inferred modules and reference Gene Ontology (GO) pathways was analyzed using the “newGeneOverlap” function from GeneOverlap R package (version 1.26.0). It utilizes Fisher exact tests to determine whether enrichment is significant (adjusted p < 0.05) and reports odds ratio and Jaccard index to denote the level of enrichment (Shen, 2014). Results are reported for the top five GO modules with the highest odds ratios. The default arguments were employed for all the analyses in this study.

### Gene Differential Expression Analysis

The “FindMarkers” function from Seurat R package (version 4.3.0) was used to identify differentially expressed genes using the Wilcoxon rank sum test. Genes with an absolute log-2-fold change larger than 1 and an adjusted p-value less than 0.05 were considered significant.

### Remaining Used Functions

Plots were created using ggplot2 R package (version 3.3.5) unless noted. igraph R package (version 4.0.0) was used for creating and manipulating graphs. The function “DotPlot” from Seurat R package (version 4.3.0) was used to visualize the gene expression with dot plot. The function “DimPlot” from Seurat R package (version 4.3.0) was used to visualize the final UMAP on a 2D scatter plot. The “EnhancedVolcano” function from EnhancedVolcano R package (version 1.18.0) was used to visualize the deferentially expressed genes. All analyses were performed using R version 4.3.0.

---

**Supplementary Figure 1:**
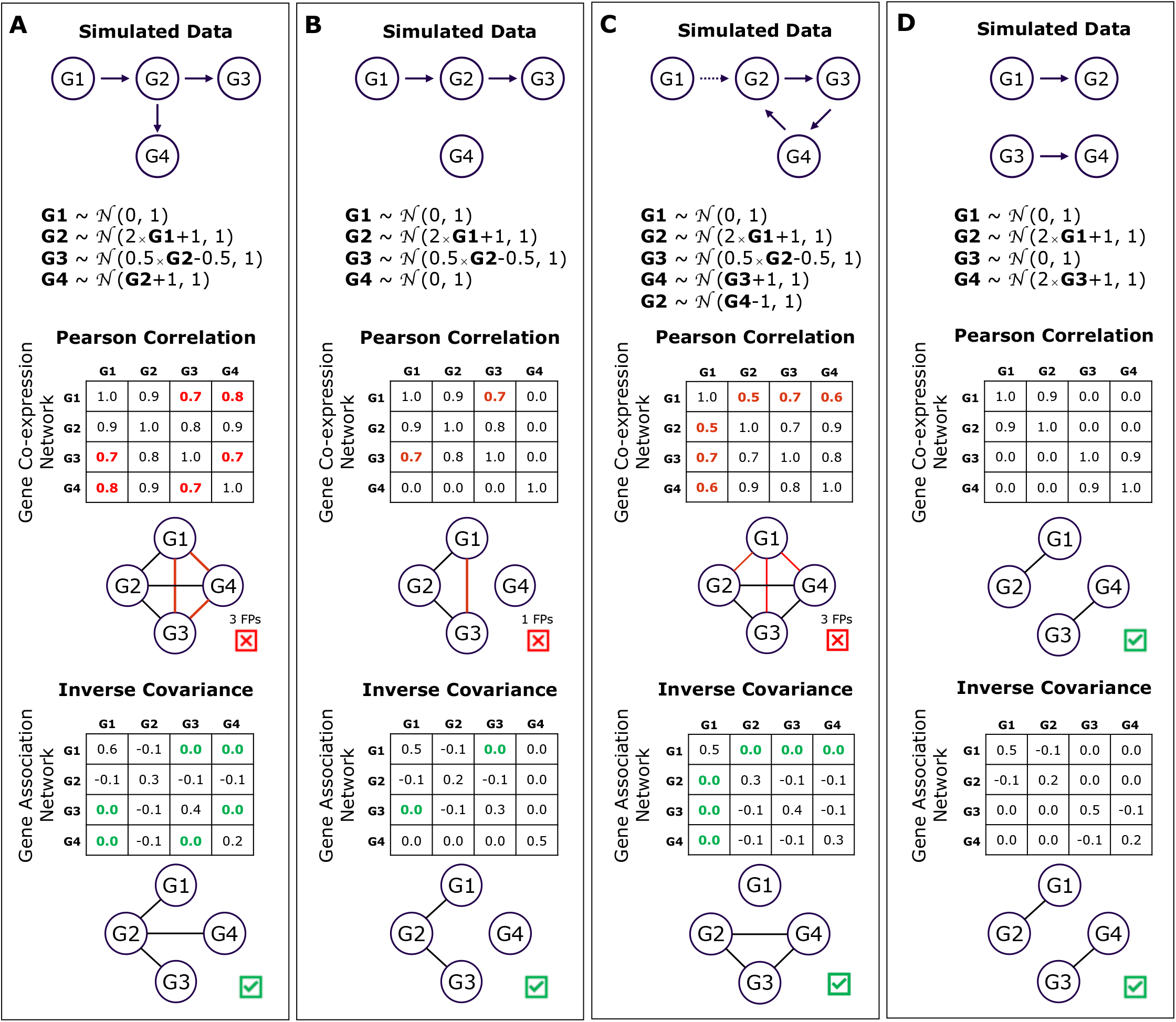
Inverse covariance matrix deciphers direct associations from indirect effects in synthetic data. **(A-D)** First row: Four unique pseudo-gene configurations representing alternative transcriptional relationships. Second row: Set of equations used to simulate pseudo-gene expressions based on predefined relationships and characteristics. Third and fourth rows: Pearson correlation matrices used for constructing gene co-expression networks, respectively. The red values in the matrices and red edges in the networks highlight indirect effects, representing false positives. Fifth and sixth rows: Inverse covariance matrices used for constructing gene association networks, respectively. The green zeros in the matrices highlight removed indirect effects.

**Supplementary Figure 2:**
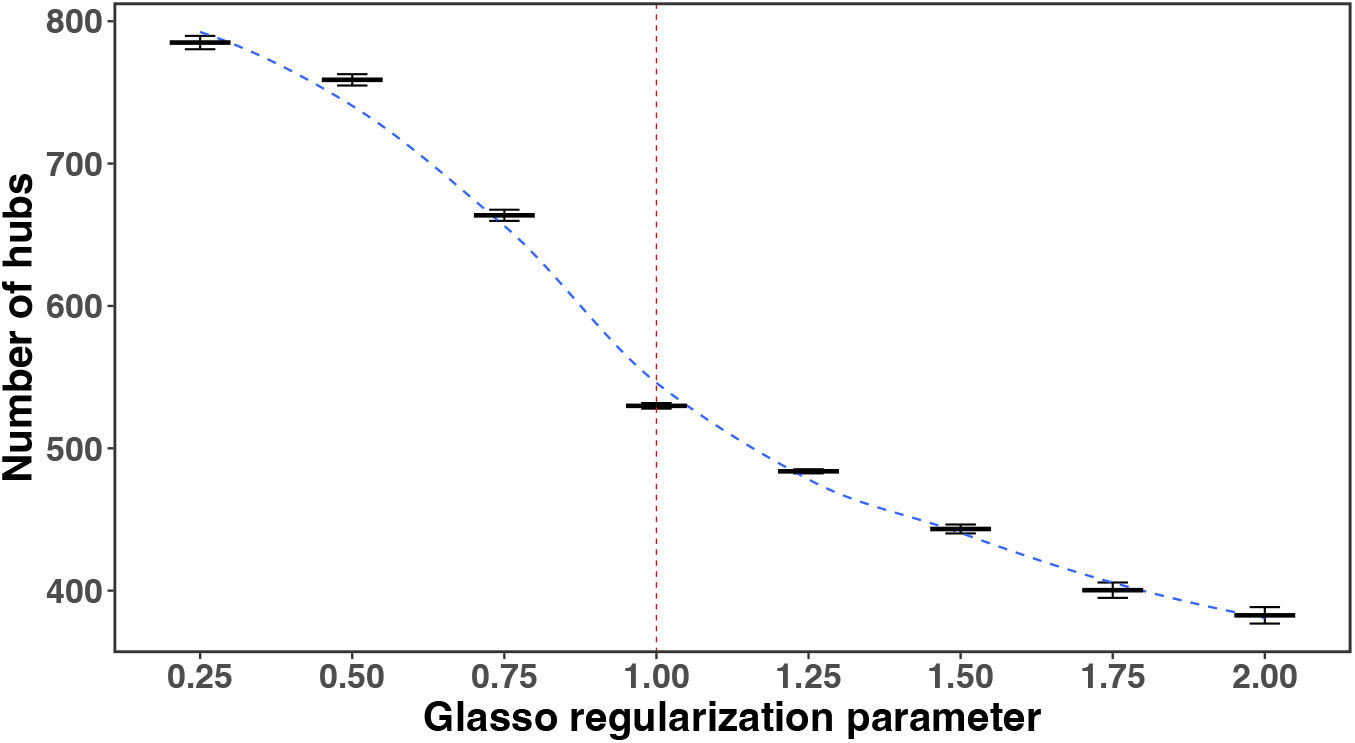
Regularization parameter selection for glasso algorithm using simulated data analysis. Six simulated scRNA-seq datasets, each comprising 5,000 cells and 10,000 genes, were used to infer gene association networks using a range of regularization parameters spanning from 0.25 to 2. The graph depicts the number of hub genes (y-axis) plotted against the regularization parameter (x-axis), incremented by 0.25. Error bars illustrate the standard deviation of hub gene counts in the inferred networks (n = 6). The blue dashed line represents the curve fitting the average number of hubs for each regularization parameter. The red dashed line signifies the selected regularization parameter of 1.

**Supplementary Figure 3:**
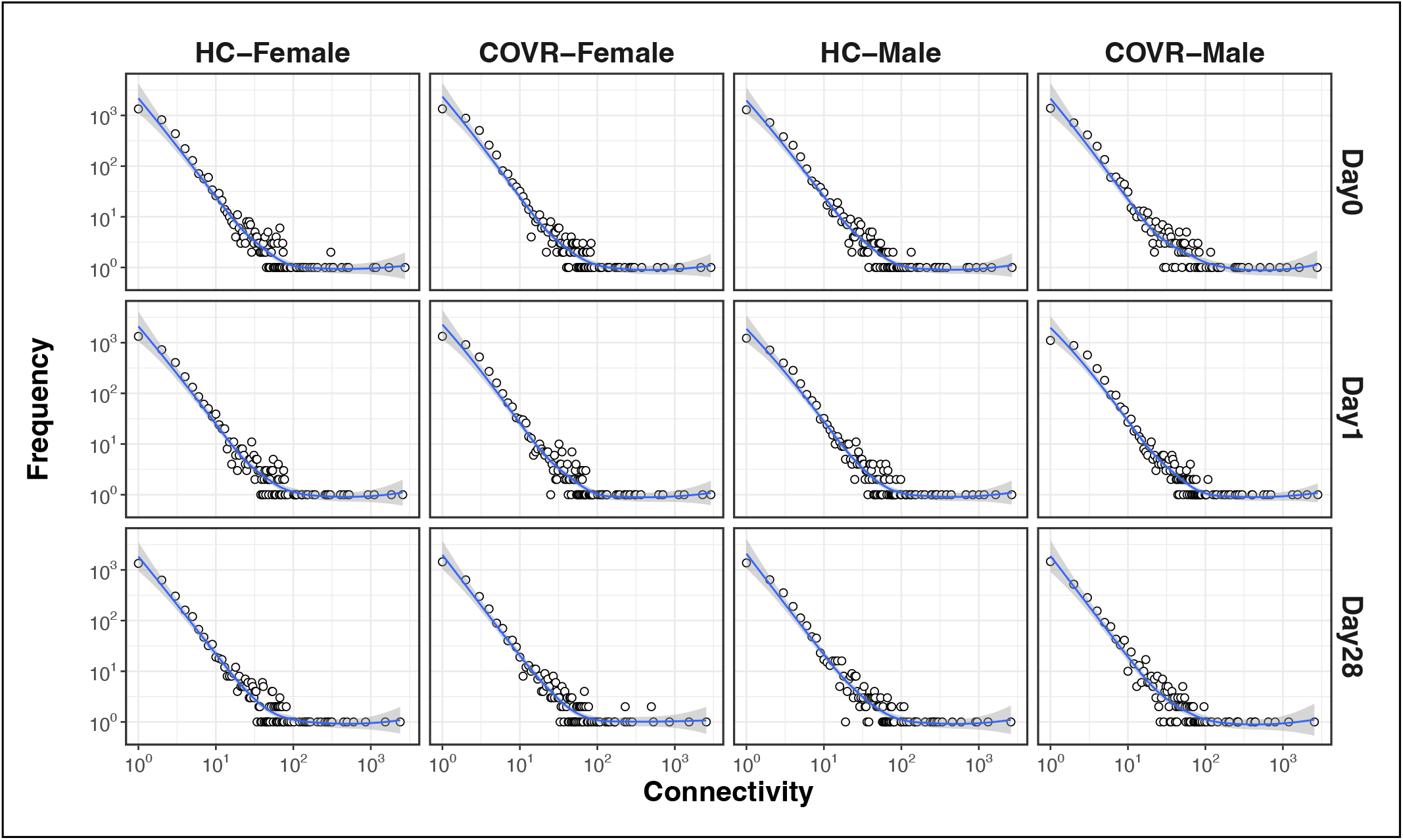
GENIX infers gene networks with heavy tail degree distribution. Illustrates the distribution of gene degrees in the networks inferred from classical monocytes in healthy control (HC) and COVID-19 recovered (COVR) female and male participants at D0, D1, and D28. The x-axis represents the degree of each gene (the number of connections), and the y-axis shows the number of genes with a specific degree.

**Supplementary Figure 4:**
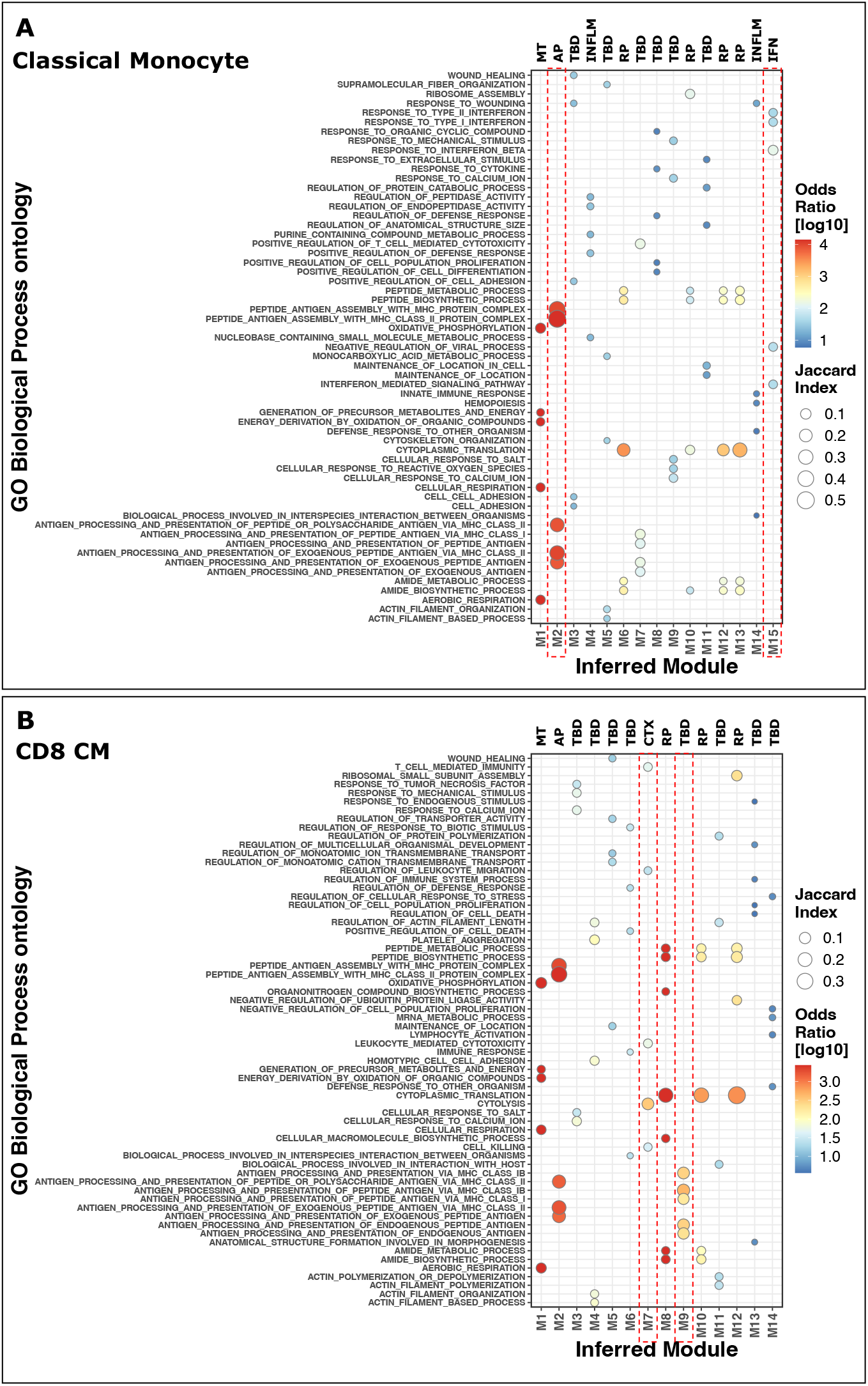
Gene enrichment analysis for annotating inferred modules functions using Gene Ontology (GO) database. The gene enrichment analysis results for classical monocytes are shown in **(A)** and for CD8 central memory cells are shown in **(B)**. The x-axis corresponds to the inferred module identifiers, while the y-axis represents the identified GO terms. The legend color-bar indicates the odds ratio, representing the strength of association between the modules and GO terms. The legend circles size displays the Jaccard index, measuring the similarity between two lists of genes. Fisher’s exact test was employed to determine statistical significance. The results are presented for the top five GO modules with the highest odds ratios and an adjusted p-value below 0.05. The top labels indicate the annotation of modules according to Figure 2C by comparing the inferred modules to the reference functional modules from (Chaussabel, 2008; Chiche, 2014). AP for antigen presenting, CTX for cytotoxic, IFN for interferon, INFLM for inflammation, MT for mitochondrial, RP for ribosomal protein, and TBD for unannotated modules. The red dashed vertical rectangles highlight modules used in the analysis of the Results section.

**Supplementary Figure 5:**
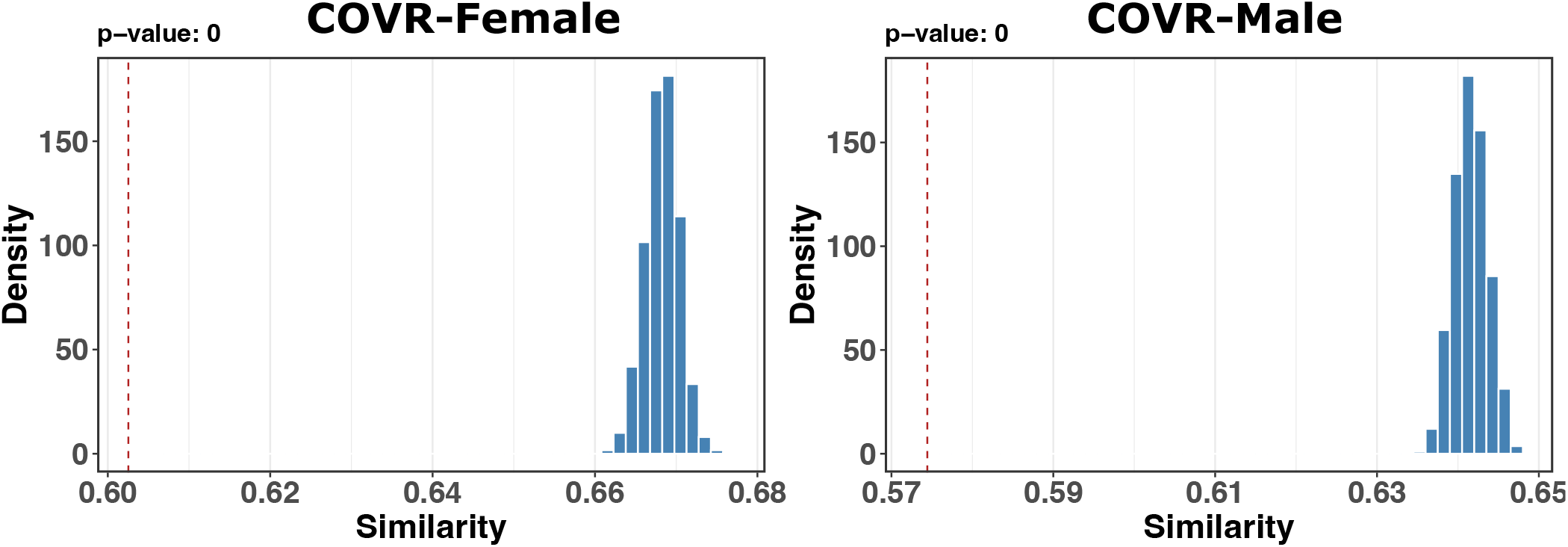
Network distinctiveness assessment using permutation test. The histogram illustrates the distribution of Jaccard index similarities (x-axis) obtained from the permutation test, comparing the pairs of networks that are sampled (n = 50,000) from a union of observed networks from classical monocytes on D0 and D1 after vaccination. The left panel depicts the result from female participants, while the right panel depicts male participants who had recovered from COVID-19 (COVR). The y-axis indicates the density of occurrences for each similarity. The vertical red dashed line denotes the observed similarity value. The displayed p-value at the top is calculated using a two-sided t-test and serves to demarcate statistically significant (p < 0.05) differences in network structures.

